# Adenomyosis and fibrosis define the morphological memory of the postpartum uterus of dairy cows previously exposed to metritis

**DOI:** 10.1101/2024.05.31.596515

**Authors:** I Sellmer Ramos, MO Caldeira, SE Poock, JGN Moraes, MC Lucy, AL Patterson

## Abstract

Optimal reproductive success following parturition in lactating dairy cows is dependent upon adequate completion of uterine involution. Failure to resolve pathogenic bacterial contamination within the first week postpartum can lead to uterine disease (metritis). Metritis is associated with decreased fertility and a failure or delay to establish pregnancy. We hypothesized that the inflammation resulting from early postpartum metritis would be associated with long-term changes in uterine morphology due to impaired uterine involution within the first 30 days postpartum (dpp). First parity Holstein cows were diagnosed with or without metritis at 7-10 dpp and uterine tissue were analyzed at 30 (Exp. 1), or 80 and 165 (Exp. 2) dpp for the presence of abnormal morphology, including abnormal invasion of endometrial glands and stroma into the myometrium (adenomyosis) using immunohistochemistry for FOXA2 (uterine gland specific marker) and presence of late postpartum endometrial fibrosis using mason’s trichrome stain (MTS). Severity of adenomyosis was determined by the number and size of adenomyotic foci, distance of foci from the endometrium-myometrium interface (EMI), and degree of fibrosis (MTS stain intensity). The presence, size, and distance from the EMI of adenomyotic foci were greater later postpartum and in cows with early postpartum diagnosis of metritis. Endometrial fibrosis was greater at the stratum basalis (at EMI) compared to the stratum compactum endometrium (near lumen) for all Exp. 2 cows, but greater endometrial fibrosis (regardless of endometrial region) was observed in cows that were diagnosed with metritis. Taken together, these data indicate that early postpartum metritis is associated with long-term modifications to the postpartum uterine morphology, including aberrant endometrial invasion into the myometrium (adenomyosis) and increased pathological fibrogenesis, leading to the presence of late postpartum endometrial fibrosis (scar tissue). Additionally, increased collagen fiber at the EMI suggests a correlation between the development of adenomyosis and fibrosis, which could possibly result from sustained endometrial inflammation caused by uterine disease.

## Introduction

A cascade of physiological, morphological, and metabolic adaptations to the transition between pregnancy, parturition and subsequent lactation characterizes the challenges faced by dairy cows after calving. Adequate completion of early postpartum events poses a significant contribution to optimal reproductive performance in lactating dairy cows during the breeding period (Lucy, 2019). Amongst these events is the process of uterine involution, which is characterized by the functional reestablishment of the uterine histoarchitecture, in order to support subsequent early embryonic development and the attachment of the conceptus to the endometrial epithelia.

Despite the success achieved by reproductive programs that target early return to cyclicity at first service, reproductive success in lactating dairy cows remains suboptimal due to factors that contribute to early embryonic loss. Amongst these factors is the failure of the endometrium to support the establishment of pregnancy, which can be attributed to simultaneous endocrine disruption and persistent local inflammation (LeBlanc, 2023).

Early postpartum uterine disease (metritis), which affects approximately 25% of the lactating dairy herd in the U.S., is caused by the invasion of pathogenic bacteria into the endometrial luminal epithelia (Pinedo et al., 2020). Failure to resolve early postpartum bacterial infection and subsequent inflammatory responses can delay or permanently impact the reestablishment of the uterine histoarchitecture in dairy cows that were previously diagnosed with metritis, and could be associated with fertility losses observed in metritic cows (Sellmer Ramos et al., 2023).

Inflammation fosters the development of benign uterine disorders, which pose implications to female fertility. Amongst these disorders is a phenomenon termed “adenomyosis”, a disorder characterized by the abnormal invasion of glandular epithelia surrounded by endometrial stroma into the muscular layer of the uterus (myometrium). Adenomyosis is an important gynecological disorder in women due to its significant impact on fertility (AlAshqar et al., 2021).

Recent studies have reported the presence of adenomyosis in dairy cattle and its positive relation with both postpartum uterine disease and endocrine disruption (Korzekwa et al., 2013; Talebkhan Garoussi et al., 2015).The mechanisms that regulate the development of adenomyosis are not completely understood, but its prevalence amongst uteri that have sustained injuries suggest a potential bridge between chronic uterine inflammation and fertility losses in dairy cows.

Thus, we hypothesized that the failure to resolve early postpartum uterine inflammation would be associated with a sustained uterine morphological memory to the disease-induced inflammation, including aberrant injury responses such as the abnormal migration of endometrial stroma and glandular epithelia into the myometrium of dairy cows that were diagnosed with metritis following parturition; and that such findings could shed light onto additional factors that can present a challenge to the successful establishment of pregnancy during the breeding period.

## Materials and Methods

### Animals and tissue collection

Study procedures were approved by the University of Missouri Institutional Animal Care and Use Committee (Protocol number 9635). For Experiments (Exp.) 1 and 2, first parity Holstein cows were selected from a large confinement herd in eastern Kansas or the University of Missouri herd. Cows with a clinical diagnosis of metritis (fetid red-brown watery vaginal discharge with a flaccid uterus) at 7-10 dpp were selected and matched with clinically healthy postpartum cows [viscous (not watery) and non-fetid discharge] that calved during the same week. For Exp. 1, Kansas herd cows were moved to the University of Missouri shortly after diagnosis, and cows from both herds were slaughtered at approximately 30 dpp [29.1 ± 1.7 dpp; mean ± SD (n = 10 metritic, n = 10 healthy)].

Cows in Exp 1. were also used for a previous study in which metritic and healthy cows were randomly assigned to either antibiotic treatment [healthy; n = 5 and metritic, n = 5; ceftiofur hydrochloride (i.m.; 2.2 mg/kg for 3 d)] or not treated (healthy, n = 5 and metritic, n = 5). Results from that experiment demonstrated no long-term effect of antibiotic treatment on a variety of study endpoints including uterine gene expression, microbiome, and inflammation (Moraes et al., 2024; Silva et al., 2024). Therefore, cows in Exp 2. were treated with ceftiofur hydrochloride at the discretion of the herdsman.

For Exp. 2, cows from the Kansas herd remained in the herd until transport to the University of Missouri 1 d prior to slaughter at approximately 80 dpp (n = 5 metritis and n = 5 control; 79.0 ± 7.5 dpp; mean ± SD) or approximately 165 dpp (n = 4 metritis and n = 5 control; 165.0 ± 4.9 dpp; mean ± SD).

Cows were slaughtered by captive bolt and exsanguination at the University of Missouri abattoir. The reproductive tract was removed, wrapped in sterile surgical drape, placed on ice in a plastic bag and transported to the laboratory. The uterine lumen was flushed with sterile saline and a cross section from the non-gravid horn that included both caruncular and intercaruncular areas was fixed in 10% neutral buffered formalin then paraffin embedded. and subjected to immunohistochemistry for FOXA2 and Masson’s trichrome staining.

### Immunohistochemistry and Mason’s Trichrome staining

For Exp. 1 and 2, immunohistochemistry for FOXA2 (marker of uterine glands) was performed on 5 μm sections according to the previously reported protocol (Sellmer Ramos et al., 2023; ab108422; 1:500 dilution; Abcam; Waltham, MA) to identify adenomyotic foci in uterine cross-sections. Foci of adenomyosis were defined by the presence of FOXA2 positive glandular epithelia surrounded by endometrial stroma within the myometrial layer of the uterus (smooth muscle). Sections were surveyed for the presence of foci using LAS X software on a Leica DM 4000 B microscope fitted with a Leica DFC 450C camera (Leica; Wetzlar, Germany) at 100X magnification. To assess severity and progression of adenomyosis, the following morphological measurements were performed per uterine section for each cow, 1) number of foci, 2) width and height mean diameter of foci, and 3) foci-to-endometrium distance. Foci diameter and distance were collected using the scale measurement tool (μm) (LAS X; Leica; Wetzlar, Germany). For Exp. 2, Masson’s trichrome staining was performed to assess endometrial fibrosis in sections within a 20 μm distance of the FOXA2-assayed sections using a kit (ab150686, Abcam; Waltham, MA) according to manufacturer instructions.

The endometrium in cross section was divided into two anatomical regions, defined as the stratum basalis (endometrial stroma region closest to the myometrium) and stratum compactum (endometrial stroma region closest to the uterine lumen). Photographs were captured of each region per section from individual cows at 200X magnification using a Leica DM 4000 B microscope fitted with a Leica DFC 450C camera (Leica; Wetzlar, Germany). The intensity of collagen fiber was quantified by calculating the mean pixel intensity of stain of collagen fibers (blue stain) using the linear measurement tool in ImageJ (NIH; Bethesda MD). To control for variance across tissue sections that can occur during the staining procedure, a background staining intensity measurement was collected from the myometrial region (smooth muscle fibers stained red), and mean intensity of collagen fiber stain was obtained by subtracting the collagen quantification from background stain.

### Statistical Analyses

Data were analyzed using SAS 9.4 (SAS Institute Inc., Cary, NC). For Exp.1 and Exp.2, the dependent variables were: 1) the number of myometrial adenomyotic foci within a uterine cross section; 2) the mean width and height diameter (μm) of adenomyotic foci; and 3) the foci-to-endometrium distance (μm). For Exp.2, an additional dependent variable was added: 4) the mean pixel intensity of stain of endometrial collagen fiber (indicative of fibrosis). Dependent variable 1 was analyzed using a negative binomial regression model that included status and days postpartum, due to the lack of a normal distribution (zero-inflated and right skewed). The dependent variables 2 and 3 were analyzed using a two-way ANOVA procedure (PROC GLM) of SAS. A model that included status (healthy or metritic) and days postpartum (30, 80 or 165 dpp) and all interactions was normally distributed and fit. Data from Exp. 1 (30 dpp) and Exp. 2 (80 or 165 dpp) were analyzed collectively to determine if early postpartum cows (30 dpp) or later postpartum cows (80 or 165 dpp) differed for number, diameter, and distance of adenomyotic foci in each respective uterine cross section. Dependent variable 4 was analyzed using a two-way ANOVA procedure (PROC GLM) of SAS, in a model that included data from Exp. 2 (80 and 165 dpp) including the fixed effects of disease status and days postpartum. Regardless of the analysis, the LSMEANS were separated by using the Tukey procedure. Significance was declared at *P* < 0.05. A statistical tendency was 0.05 < *P* < 0.10. Data are presented as lsmeans ± SEM unless stated otherwise.

## Results

Using FOXA2 expression (**Fig. 1 A**), a marker of uterine glands, and masons trichrome staining (MTS; **Fig. 1 B**) adenomyotic foci were identified in 7 cows (35%) at 30 dpp, 8 cows (80%) at 80 dpp, and 7 cows (78%) at 150 dpp. When stratifying by disease status (healthy or metritic, **Fig. 1C-J**), for Exp 1, at 30 dpp presence of adenomyosis was evenly distributed between healthy (4/10, **Fig. 1 C, E**) and metritic cows (3/10, **Fig. 1 G, I**). However, for Exp 2, all metritic cows had adenomyosis (80 dpp, 5/5; 165 dpp, 5/5; **Fig. 1 H, J**), whereas approximately half of the healthy cows had adenomyosis (80 dpp, 3/5; 165 dpp, 2/4; **Fig. 1 D, F**).

**Figure 1.**
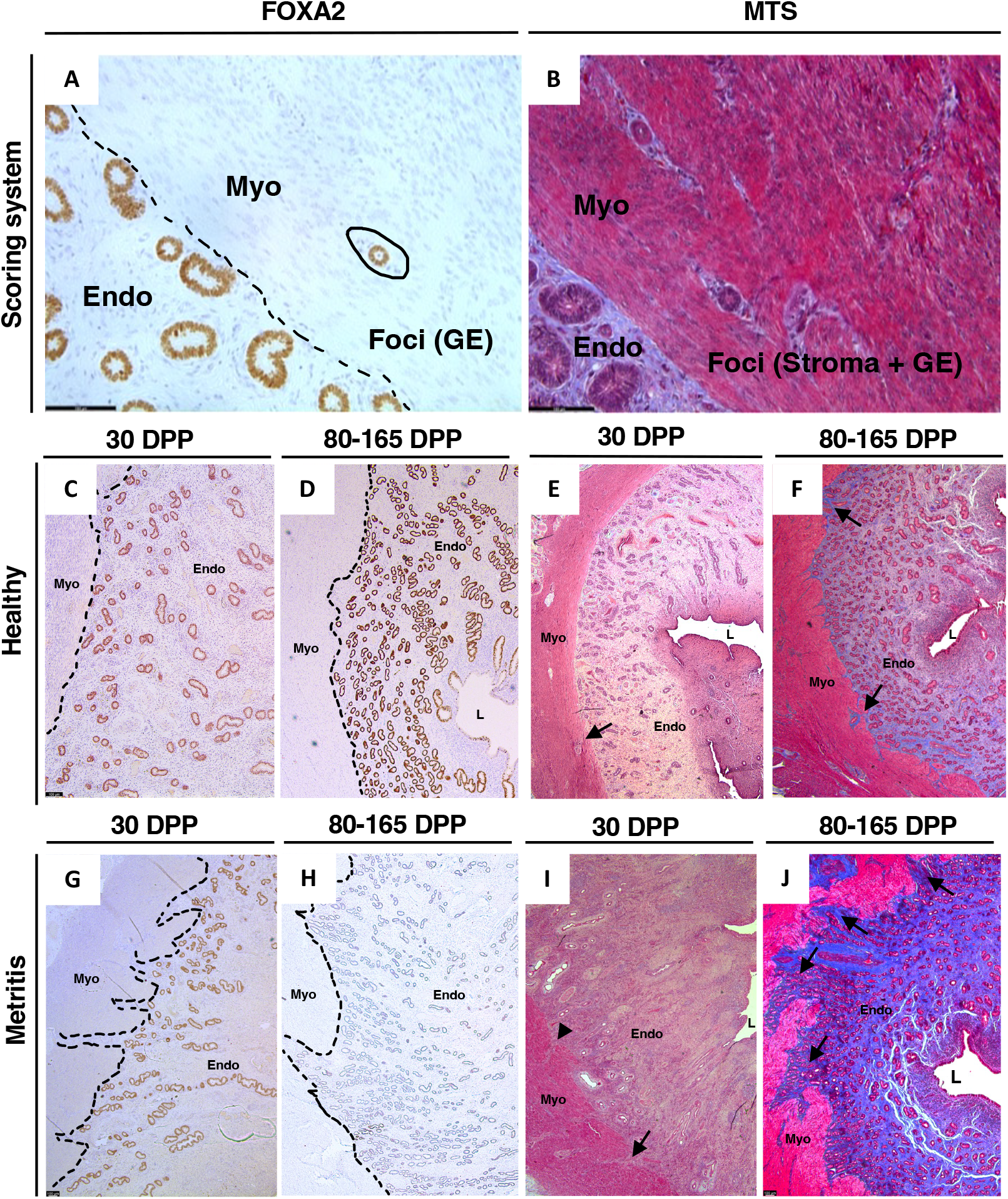
Uterine histological cross sections of healthy (C, D, E, F) and metritic (G, H, I, J) cows at early (Exp. 1, 30 dpp; C, G, E, I) and late (Exp. 2, 80 and 165 dpp; D, F, H, J) days postpartum (dpp) using FOXA2 (gland marker; A, C, D, G, H) and Mason’s Trichrome Stain (MTS; connective tissue stain; B, E, F, I, J) as a method to identify the presence of adenomyotic lesions in the myometrium (40X magnification; A, B) and to assess the severity of myometrial invasion by endometrial stroma and glandular epithelium (2.5X magnification; C-J). Myo = myometrium; Endo = endometrium; L = uterine lumen; GE = glandular epithelium; Foci = adenomyotic foci. Bar = 100 μm.

There was an effect of days postpartum on number of adenomyotic foci per uterine cross section and an effect of days postpartum on the diameter of adenomyotic foci (**Table 1**). The total number of adenomyotic foci per uterine cross section was greater in cows at 165 dpp or 80 dpp (1.484 ± 0.35 and 1.409 ± 0.34, respectively) when compared to 30 dpp (0.141 ± 0.32) (*P* < 0.0001). When assessing adenomyotic foci diameter, the greatest diameter was found in cows at 80 dpp (352.68 ± 28.6) compared with 165 dpp (237.96 ± 32.8) or 30 dpp (228.05 ± 32.0) (*P =* 0.0085).

**Table 1.** Summary of least squares means for adenomyosis foci diameter, foci-to-endometrium distance and foci number for variables days postpartum, disease status, and all significant two-way interactions.

| Variable | LSM | SE | 95% Confidence Interval |  |
| --- | --- | --- | --- | --- |
|  |  |  | Lower Bound | Upper Bound |
| Disease status |  |  |  |  |
| Healthy |  |  |  |  |
| Foci diameter | 220.43<sup>a</sup> | 28.232 | 165.10 | 275.76 |
| Foci-to-endometrium distance | 205.73<sup>a</sup> | 40.922 | 125.52 | 285.94 |
| Foci number per uterine cross section | 0.57 | 0.296 | -0.02 | 1.15 |
| Metritic |  |  |  |  |
| Foci diameter | 325.36<sup>b</sup> | 22.383 | 281.49 | 369.23 |
| Foci-to-endometrium distance | 383.62<sup>b</sup> | 32.444 | 320.03 | 447.21 |
| Foci number per uterine cross section | 1.25 | 0.355 | 0.56 | 1.95 |
| Days postpartum |  |  |  |  |
| 30 |  |  |  |  |
| Foci diameter | 228.05<sup>a</sup> | 32.012 | 165.31 | 290.79 |
| Foci-to-endometrium distance | 264.86 | 46.401 | 173.91 | 355.81 |
| Foci number per uterine cross section | 0.14<sup>a</sup> | 0.322 | -0.49 | 0.77 |
| 80 |  |  |  |  |
| Foci diameter | 352.68<sup>b</sup> | 28.632 | 296.56 | 408.80 |
| Foci-to-endometrium distance | 334.07 | 41.503 | 252.72 | 415.42 |
| Foci number per uterine cross section | 1.41<sup>b</sup> | 0.336 | 0.75 | 2.07 |
| 165 |  |  |  |  |
| Foci diameter | 237.96<sup>a</sup> | 32.802 | 173.67 | 302.25 |
| Foci-to-endometrium distance | 285.11 | 47.548 | 191.92 | 378.30 |
| Foci number per uterine cross section | 1.48<sup>b</sup> | 0.351 | 0.79 | 2.17 |

<sup>a,b</sup>Means with differing superscripts are significantly different at $P < 0.05$

There was an effect of disease status (healthy or metritic) on the andenomyotic foci-to-endometrium distance and the diameter of adenomyotic foci per uterine cross section (*P* < 0.05) (**Table 1**). The distance of foci within the myometrium to the endometrial border was greater in cows that were diagnosed with metritis (383.6 ± 32.4) when compared to cows that were deemed healthy (205.7 ± 40.9) at diagnosis (7 to 10 dpp) (*P* = 0.0009). Similarly, cows that were deemed metritic had a greater mean foci diameter (325.4 ± 22.4) when compared to cows that were healthy (223 ± 41.0) (*P* = 0.004). Lastly, there was a tendency (*P* = 0.077) for a greater number of adenomyotic foci per uterine section in diseased (metritic; 1.25 ± 0.25) compared to healthy (0.56 ± 0.3) cows. This suggests an association between early postpartum metritis with the severity of adenomyosis (foci-to-endometrium distance, foci diameter, and number of foci).

For Exp. 2 (80 and 165 dpp), there was an effect of both disease status and endometrial layer independently on the intensity of endometrial fibrosis (*P* < 0.0001, **Table 1**). The mean pixel intensity of blue collagen fiber stain was greater in the deep endometrial stromal layer [(near the myometrial border, **Fig. 2 B, E**) 107.6 ± 3.28] when compared to the superficial endometrial stromal layer [(near the luminal epithelium, **Fig. 2 C, F**) 72.5 ± 3.28, *P* < 0.0001], and was greater in the endometrium of cows that were diagnosed with early postpartum metritis (103.1 ± 3.19; **Fig. 2 D**) when compared to cows that were deemed healthy (76.96 ± 3.39; *P* < 0.0001; **Fig. 2 A**) at both 80 and 165 dpp. Of note, there was little-to-no detectable levels of collagen fiber stain at 30 dpp regardless of status (healthy or metritic) (see **Fig. 1 E, I**; quantified results not presented). There was also an interaction between disease status and layer (*P* < 0.05), such that the greatest intensity of collagen fiber stain was observed in the deep endometrial layer of cows that had uterine disease (127.95 ± 4.51) when compared to the same layer of cows that were healthy (87.22 ± 4.77) (*P* < 0.0001). Taken together, these data suggest a correlation between the extent of fibrosis and the abnormal invasion of endometrial stroma and uterine glands into the myometrial layer of the uterus and subsequent development of adenomyosis.

**Figure 2.**
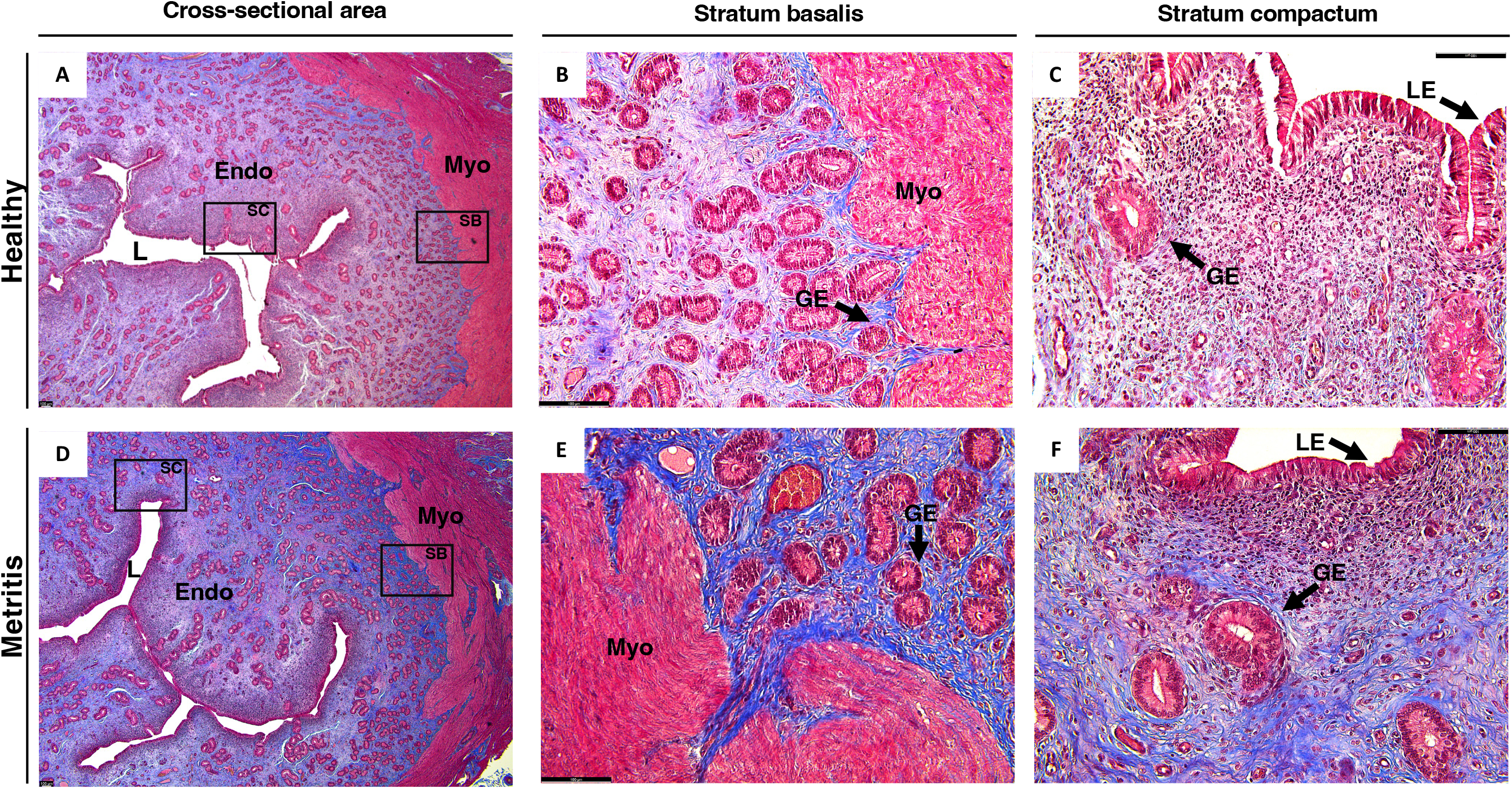
Endometrial cross sections of late postpartum (Exp. 2; 80 and 165 dpp) cows stained with Mason’s Trichrome to detect and quantify collagen fiber (blue stain) within the stratum basalis [SB; endometrial stroma closest to the myometrium (Myo); B, E] and stratum compactum [SC; endometrial stroma closest to the luminal epithelium (LE); C, F] layers of the endometrium (Endo) of healthy (A, B, C) and metritic (D, E, F) cows as a method to assess the severity of postpartum uterine fibrosis. Insets in A and D correspond to images B & C and E & F, respectively. Myo = myometrium (red stain); L = uterine lumen; GE = glandular epithelium; scale bar = 100 μm.

## Discussion

This series of observations demonstrates significant evidence for the negative effect that early postpartum uterine disease poses to the reestablishment of normal uterine histoarchitecture. We observed that the uterus of cows diagnosed with metritis between 7 to 10 dpp had increased evidence of abnormal invasion of endometrial glands into the myometrium (adenomyosis). Although unclear, the pathogenesis of adenomyosis is hypothesized to be initiated by the disruption of the endometrial-myometrial border, followed by the subsequent invasion of endometrial tissue (glandular epithelia surrounded by stromal fibroblasts) into the myometrium (Bulun et al., 2021; Li et al., 2024). Local trauma at the endometrial-myometrial interface (*e.g*., parturition, uterine infection) associated with myometrial hyperperistalsis leads to the activation of the tissue injury-repair mechanism (TIAR), which is marked by pro-inflammatory signaling (Leyendecker et al., 2009; Scull et al., 2010). Chronic endometrial inflammation, particularly when induced by secreted cytokine activity, has been demonstrated to promote endometrial infiltration and migration (Bodur et al., 2015). Accordingly, we demonstrated in a series of *in vitro* studies that in response to LPS treatment (modulator of pro-inflammatory responses), 80 and 165 dpp-derived bovine endometrial epithelial cells (bEECs) upregulated of the expression of pro-inflammatory markers, such as TNF-a, CXCL8, PTGS2 and SAA3 (Caldeira et al., 2022), and genes involved in extracellular matrix (ECM) organization and cell-cell adhesion, including *COL1A1, COL1A2, MMP2, MMP9* and *TGFBI* (Caldeira et al., 2023). These novel findings describe the first evidence of an epithelial immunological memory within the uterus of lactating dairy cows following a diagnosis of metritis and association with the abnormal invasion of endometrial glands and stroma into the myometrium.

Herein we found greater fibrosis (collagen fiber staining) in the endometrium of later postpartum cows (Exp. 2, 80 and 165 dpp) compared with the cows within the time of expected completion of uterine involution (early postpartum, 30 dpp). In the context of TIAR, fibrogenesis (development of fibrosis) is a biological process that results from TIAR-induced inflammation, which consequently induces ECM synthesis and deposition upon fibroblast activation (Wynn, 2007). Recent studies investigating the uteri of women that previously sustained endometrial injury have found a link between adenomyotic lesions and fibrosis. Chronic myometrial hyperperistalsis led by insult to the endometrial-myometrial interface has been associated to muscle fiber disruption in focal adenomyotic areas, which subsequently triggers a repeated tissue injury and repair response (Guo and Groothuis, 2018; Kobayashi et al., 2021) resulting in epithelial-to-mesenchymal transition, fibroblast-to-myofibroblast transdifferentiation, and subsequent fibrogenesis (Liu et al., 2016; Xiao et al., 2020; Yildiz et al., 2023). Moreover, the presence of fibroblast subtypes with aberrant gene expression was identified in a recent single-cell RNA-seq study that aimed to characterize the cellular landscape of adenomyotic lesions of women (Niu et al., 2023). Accordingly, we observed that the intensity of fibrosis in later postpartum cows (Exp. 2, 80 and 165 dpp) was increased in cows that were diagnosed with metritis, and more evidently in the basal (deep) endometrium nearest the myometrium. In addition to fibroblast differentiation, TIAR-induced fibrogenesis can be exacerbated by the presence of chronic inflammation, more notably attributed to the presence of macrophages and regulatory T cells (Bacci et al., 2009)(Bacci et al., 2009; Xiao et al., 2020). Based on joint metagenomics and global endometrial transcriptome, we recently demonstrated that the early postpartum metritis-microbiome of Exp. 2 (80 and 165 dpp) cows modulated the endometrial transcriptome at mid-lactation (breeding period) and suggested the presence of residual immune cells within the uterus of cows that were diagnosed with early postpartum uterine disease (Silva et al., 2024). Such findings suggest a timeline for the development of fibrotic-associated endometrial invasion and could explain why little-to-no fibrosis was observed in Exp. 1 (30 dpp) cows.

Taken together, our data brings novel insights to the multimodal effect of early postpartum uterine disease on the fertility of lactating dairy cows. We demonstrated that the combination of tissue injury and inflammation, as a result of early postpartum bacterial invasion into the endometrium, impaired proper tissue remodeling upon uterine involution (Exp.,1 30 dpp) and demonstrated long-term modifications to the normal uterine histoarchitecture (Exp. 2, 80 and 165 dpp), suggesting that metritis may modulate the reestablishment of the functional uterine landscape due to the presence of a morphological memory, characterized by chronic inflammation, abnormal endometrial migration and the presence of non-functional tissue (fibrosis). We hypothesize that the presence of a morphological memory to factors that negatively impact tissue repair could play a significant role in the fertility of dairy cows at the uterine level and requires further investigation.

